# Scrublet: computational identification of cell doublets in single-cell transcriptomic data

**DOI:** 10.1101/357368

**Authors:** Samuel L. Wolock, Romain Lopez, Allon M. Klein

## Abstract

Single-cell RNA-sequencing has become a widely used, powerful approach for studying cell populations. However, these methods often generate multiplet artifacts, where two or more cells receive the same barcode, resulting in a hybrid transcriptome. In most experiments, multiplets account for several percent of transcriptomes and can confound downstream data analysis. Here, we present Scrublet (<u>S</u>ingle-<u>C</u>ell <u>R</u>emover of Do<u>ublet</u>s), a framework for predicting the impact of multiplets in a given analysis and identifying problematic multiplets. Scrublet avoids the need for expert knowledge or cell clustering by simulating multiplets from the data and building a nearest neighbor classifier. To demonstrate the utility of this approach, we test Scrublet on several datasets that include independent knowledge of cell multiplets.

## Introduction

Single-cell RNA-sequencing (scRNA-seq) is a powerful and accessible approach for studying complex biological systems. It is quickly becoming a standard tool for unbiased characterization of tissue cell types and high-resolution reconstruction of differentiation trajectories [1]. Droplet microfluidic [2–4] and well-based [5–8] technologies now enable the relatively inexpensive, high-throughput isolation and barcoding of cell transcriptomes. However, these methods suffer from the problem of cell multiplets, where a mixture of two or more cells is reported as a single cell in the data.

Most scRNA-seq technologies co-encapsulate cells and barcoded primers in a small reaction volume (droplets or wells), thereby associating the mRNA of each cell with a unique DNA barcode. Multiplets arise when two or more cells are captured within the same reaction, generating a hybrid transcriptome (**Fig. 1A**). Cell multiplets are a concern when interpreting the outcome of scRNA-seq experiments, because they suggest the existence of intermediate cell states that may not actually exist in the sample. Such artifactual states can confound downstream analyses by appearing as distinct cell types, bridging cell states, or interfering in differential gene expression tests and inference of gene regulatory networks (**Fig. 1B**).

**Figure 1.**
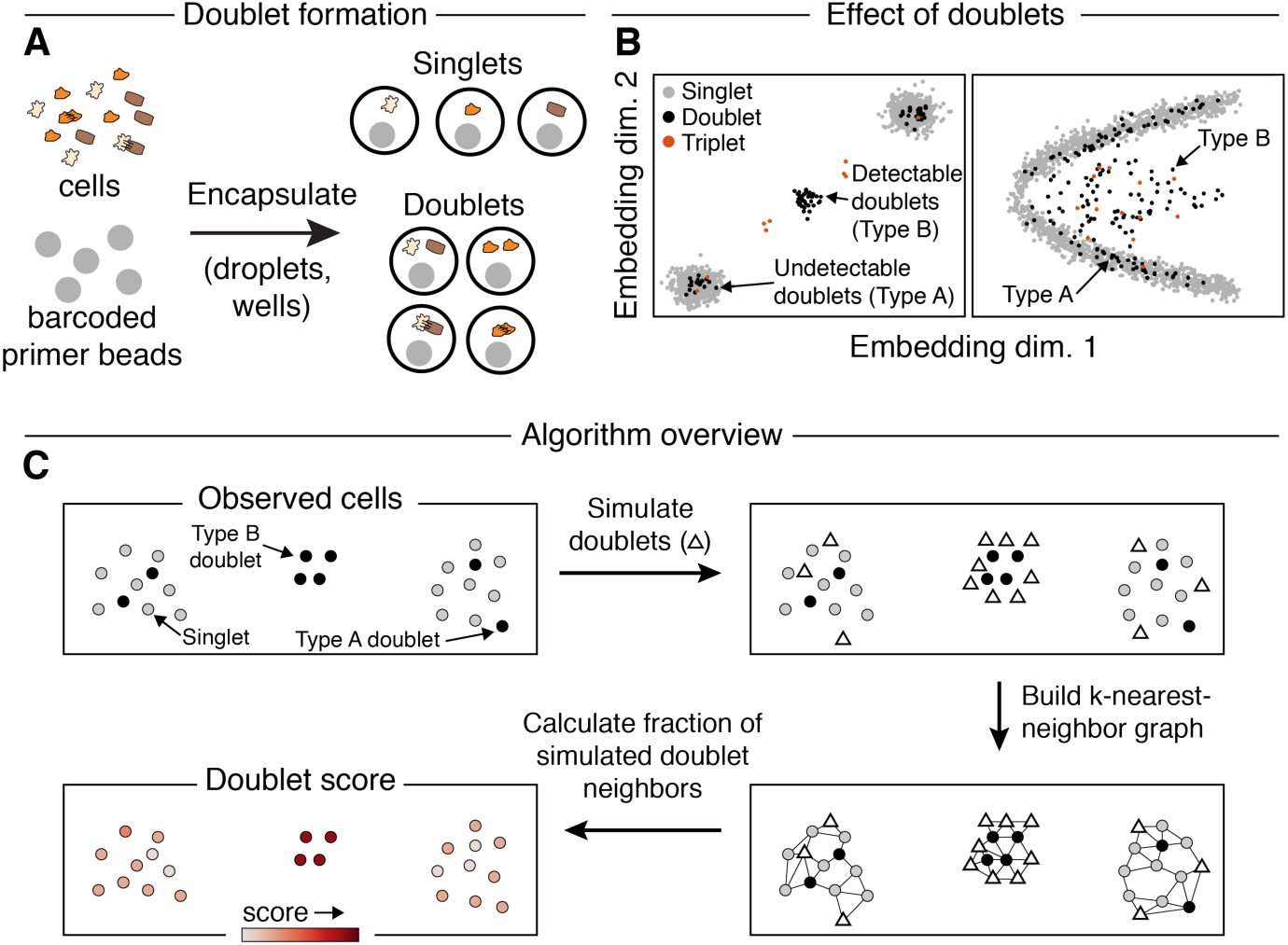
A computational approach for identifying doublets in single-cell RNA-seq data. **(A)** Schematic of doublet formation. Multiple cells are co-encapsulated with a single barcoded bead, either randomly or as aggregates, resulting in the generation of a hybrid transcriptome. **(B)** Multiplets involving highly similar cells (“Type A”) may be difficult to distinguish from single cells, while multiplets of dissimilar cells (“Type B”) generate qualitatively new features, such as distinct clusters (left) or bridges (right). **(C)** Overview of the Scrublet algorithm. Doublets are simulated by randomly sampling and combining observed cells, and the local density of simulated doublets, as measured by a nearest neighbor graph, is used to calculate a doublet score for each observed cell.

In a typical scRNA-seq experiment, at least several percent of all capture events are multiplets [2–5]. Multiplets can form as a result of cell aggregates or through random co-encapsulation of more than one cell per droplet or well. The rate of random co-encapsulation can be reduced by processing very dilute cell suspensions. However, in practice it is often favorable to work with high cell concentrations in order to capture a large number of cells within a short amount of time and to reduce reagent costs. Additionally, multiplets resulting from cell aggregates cannot be eliminated by simply reducing cell concentration. Pre-sorting cells into wells can overcome these problems [9, 10], but at a cost in throughput. Thus, rather than avoiding multiplets, it would be useful to identify them, either computationally or through experimental means.

## The case for a computational approach to multiplet inference

Ideally, one would identify multiplet events experimentally through appropriate assay designs. At the time of writing, we noted five existing experimental strategies for multiplet detection, summarized in **Table 1**. However, none of the existing methods can yet be implemented routinely for all scRNA-seq experimental designs (see “Limitations” in **Table 1**). It would therefore be useful to have a computational strategy to infer the identity of multiplets directly from data.

**Table 1.** Experimental methods for multiplet detection.

| Method name and references | Approach | Limitations |
| --- | --- | --- |
| *Species mixing*<br>[2, 3] | Cells from different species (e.g., mouse and human) are mixed and barcoded. Multiplets are detected as cell barcodes associated with transcripts from both species. Assuming 1:1 mixing, the identified multiplets represent half of all multiplets, as the remaining half are intra-species multiplets. | Measures multiplet rate but does not facilitate detection of multiplet cell states in typical experimental samples from a single organism |
| *Natural genetic variation*<br>[11] | By mixing together cells from comparable samples from multiple genotyped individuals, genetic variants in transcripts can be used to assign each cell barcode to one individual, or in the case of multiplets, to multiple individuals. Only inter-individual multiplets can be identified, so the fraction of detectable multiplets increases with the number of individuals. | Limited to samples with high genetic diversity<br>Only possible if samples from different individuals can be pooled and assayed simultaneously |
| *Genetic labeling*<br>[12-14] | Unique, expressed, genetic labels are introduced into the cell sample prior to collection. Multiplets can then be detected as cell barcodes with multiple distinct genetic labels. | Introduction of genetic labels is currently possible only for cultured cells or limited *in vivo* conditions<br>Labeling may perturb the cells |
| *Cell “hashing”*<br>[15, 16] | Cells are split into multiple wells, and each is labeled with sample-specific oligonucleotide tags, using antibodies or chemical approaches. Samples are then pooled prior to scRNA-seq. Multiplets are identified as cell barcodes associated with multiple oligo sequences. | Not well suited for very small or fragile samples that cannot be split and recombined |
| *Cell encapsulation at multiple cell concentrations* | After processing the same input sample at multiple cell concentrations, multiplet-specific cell states can be detected by finding cell states whose proportion increases with the cell concentration. | Requires at least two runs for each sample<br>Requires sufficient cells |

Until now, two simple computational methods have been implemented to exclude putative multiplets: (1) exclude cell barcodes with unusually high numbers of detected transcripts; and (2) manually curate data, excluding cell clusters that co-express marker genes of distinct cell types [1]. Both of these methods have drawbacks. As we will show later, the former method often performs poorly because it assumes that cells contain similar amounts of RNA, when in reality samples with diverse cell types or cells in different cell cycle stages are expected to have a wide range in the number of transcripts per cell. The latter method requires expert knowledge and careful annotation of the data. Below, we propose a computational approach, Scrublet (<u>S</u>ingle-<u>C</u>ell <u>R</u>emover of Do<u>ublet</u>s), for identifying multiplets and apply the method to several datasets that include some measure of ground truth labels for cell multiplets.

Briefly, our method involves two steps. First, doublets (multiplets of just two cells) are simulated from the data by combining random pairs of observed transcriptomes. Second, each observed transcriptome is scored based on the relative densities of simulated doublets and observed transcriptomes in its vicinity. Because doublets formed by cells with divergent expression profiles may be easier to detect and have more significant consequences on downstream analyses than those formed by similar cells, we incorporate this distinction into Scrublet by predicting the fraction of doublets that belong to each class. The next section discusses these two classes of doublets in greater detail.

## Defining Type A and Type B multiplet-associated errors

Multiplets can have varying consequences for downstream analyses, depending on, in part, whether they arise from averaged measurements of cells of the same or different types (**Fig. 1B**). We accordingly define two classes of multiplet-associated errors:

*“Type A” errors:* multiplets arising from combination of cells that are similar in gene expression. These are expected to result in quantitative changes in the gene expression and abundance of a cell cluster that is otherwise dominated by singlets (i.e., transcriptomes of single cells). We would expect the impact of Type A errors to be small if multiplet events are rare, because multiplets that become embedded in a manifold already dense with singlets will have little effect on gene expression or population abundance estimates.

*“Type B” errors:* multiplets arising from combination of cells with distinct gene expression. Type B errors generate new features in single-cell gene expression data, such as clusters, “branches” from an existing cluster, or “bridges” between clusters, and thus are more likely to lead to qualitatively incorrect inferences from the data.

In practice, the degree to which multiplets can be cleanly associated with these two categories will depend on the precise structure of the single-cell manifold, so the classification should be taken as a functional distinction with respect to a specific manifold construction approach used in data analysis. For example, a multiplet state might be absorbed within a large singlet cluster in one analysis (creating a Type A error) but be detectable as a separate structure in another (Type B error). Therefore, a doublet detection method capable of both predicting doublets and estimating the Type B error rate for a given analysis method would be a powerful tool.

## Method

The approach we developed, Scrublet, focuses on Type B errors. It estimates the fraction of multiplets that are predicted to generate Type B errors and offers a method to identify and remove these multiplets. We restrict ourselves specifically to doublets, since these make up >97% of multiplets in an experiment with a <5% multiplet rate with full cell dissociation. However, in principle the approach could be readily extended to higher-order multiplets.

Our method is motivated by three assumptions. First, we assume that gene expression space is high-dimensional and sparsely populated by cells, such that that doublets between cells of two distinct types will likely fall into an otherwise unoccupied region of gene expression space. Second, we assume that among all observed transcriptomes, multiplets are relatively rare events. The third assumption is that all cell states contributing to doublets are also present as single cells elsewhere in the data. Conditions under which these assumptions might be invalidated are considered in the **Discussion**.

With these assumptions, putative Type B doublets can be identified through the following steps (**Fig. 1C**):

1. Generate “simulated doublets” through linear combination of pairs of randomly sampled observed cell transcriptomes.
2. Merge observed transcriptomes (which include yet-unknown doublets) and simulated doublets and embed on a single-cell state manifold.
3. For each observed transcriptome *i* or simulated doublet *i^′^*, define the doublet score *f_i_*, *f_i_^′^*, as the abundance ratio of simulated doublets to observed transcriptomes in the neighborhood of *i* or *i^′^* on the cell state manifold.
4. Set a doublet score threshold, *θ*, based on the bimodal distribution of *f_i_^′^*. A bimodal distribution of *f_i_^′^* arises because rare Type B doublets will have a significantly higher fraction of simulated doublet neighbors than individual cells or Type A doublets, which are surrounded by a higher density of true single cells. Simulated doublets with *f_i_^′^* < *θ* correspond to Type A doublets and those with *f_i_^′^* > *θ* to Type B doublets.
5. Calculate the “detectable doublet fraction”, defined as the fraction of simulated doublets with *f_i_^′^* > *θ*. *ϕ*_*D*_ is an estimator for the fraction of observed doublets to generate Type B errors with respect to the chosen embedding.
6. Classify observed transcriptomes with *f_i_* > as putative Type B doublets.

In the **Extended Methods**, we present a more detailed description of our algorithm, including a discussion of setting the doublet score threshold.

Our strategy avoids the need to cluster data or predefine cell state marker genes and belongs to a broader class of “target-decoy” classification methods used to filter poor quality data [17]. As with other such methods, it is useful, though not always necessary, to have an independent expectation for the error rate (here, the doublet rate estimated during sample collection).

In our specific implementation of this approach, we construct a low-dimensional embedding (Step 2 above) by applying principal component analysis (PCA) to the observed transcriptomes and simulated doublets. We then build a k-nearest-neighbor (kNN) graph to measure the density of simulated doublets in the vicinity of each cell (Step 3), calculating the doublet score for each transcriptome as the fraction of its *k* neighbors that are simulated doublets (**Fig. 1C**). This implementation is suitable for routine use, with classification of datasets of tens of thousands of cells requiring only a few minutes.

## Results

The results are organized into five sections. First, we test Scrublet on simulated datasets in order to assess its performance and limitations under simplified conditions where there is perfect knowledge of singlet and doublet identity. We then apply Scrublet to three experimental datasets, each of which provides some form of independent “ground truth” for doublet identity. Finally, we apply Scrublet to our own recently published hematopoiesis dataset, which presents a complex continuum of well-characterized cell states and where doublets can be identified through prior knowledge.

### Performance on simulated data

Using pedagogical tests on simulated data, our goal is to demonstrate that (a) it is possible to use the proposed approach as a classification scheme; and (b) the detectable doublet fraction, *ϕD*, can be used to estimate the sensitivity of the classifier, i.e., the fraction of true doublets that one might be able to identify using this approach alone.

Using the Splatter package [18], we simulated single-cell data in the form of distinct cell clusters or as a continuum of cell states (**Fig. 2A**). Varying the number and size of simulated cell clusters, the doublet detector accurately identified up to 99% of doublets that were generated between cells from different clusters (Type B doublets) with 99% precision, but only if clusters were sufficiently well-separated (**Fig. 2B,C**). For poorly separated groups of cells that did not form distinct clusters, the recall dropped below 10%. As expected, doublets formed by cells from within the same cluster (Type A doublets) were virtually indistinguishable from singlets using our method. However, Scrublet’s estimate for the detectable doublet fraction (*ϕD*), i.e., the fraction of simulated doublets above the doublet score threshold, accurately predicted the recall, suggesting that it serves as a useful tool for measuring the impact of doublets in a given analysis (**Fig. 2E,F**).

**Figure 2.**
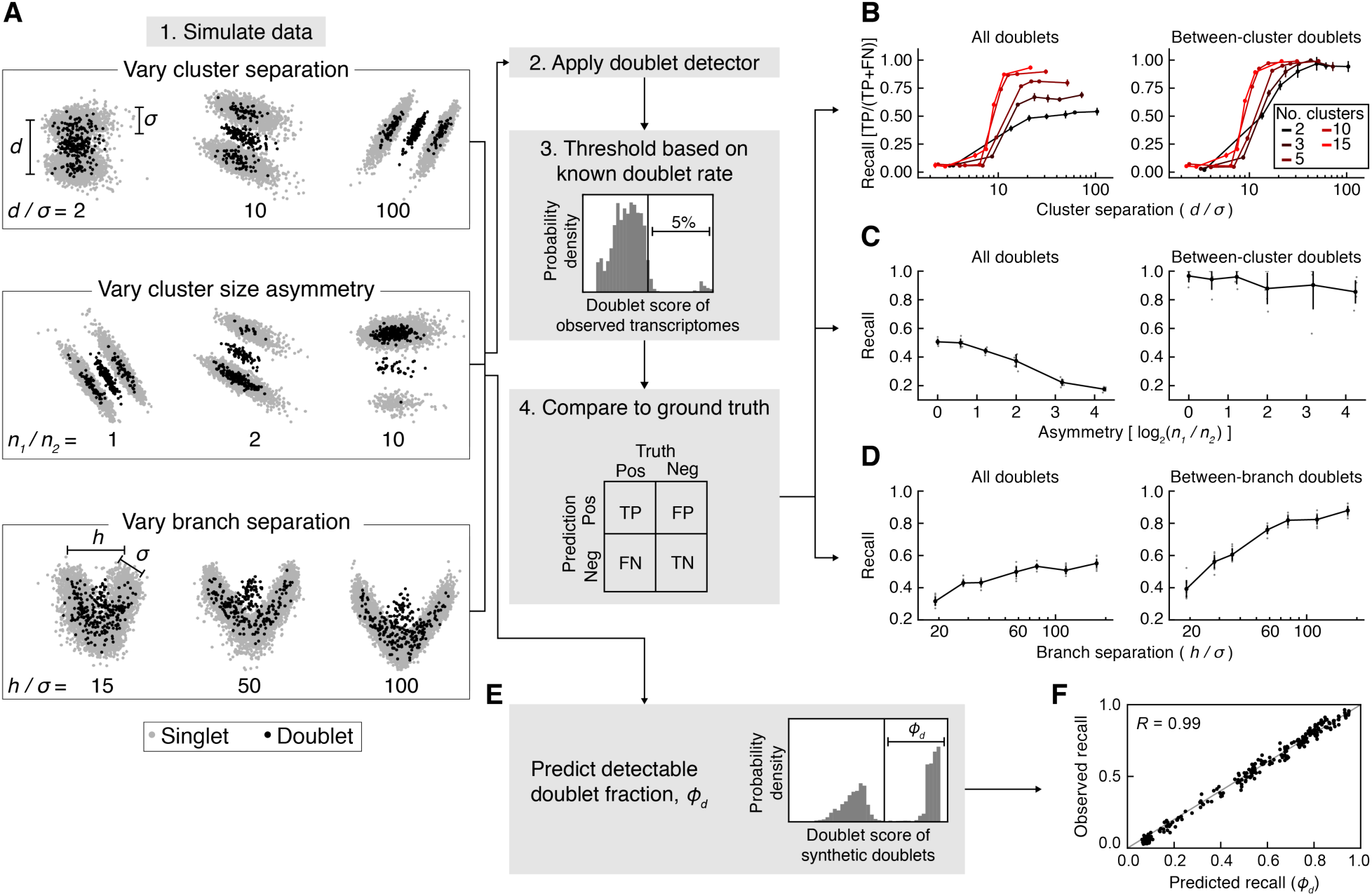
Application of Scrublet to simulated data. **(A)** Schematic summary of simulations for testing Scrublet. *d*, inter-cluster variance; *σ*, intra-cluster variance; *n_1_*, size of larger cluster; *n_2_*, size of smaller cluster; *h*, inter-branch variance. **(B)** Evaluation of doublet detector performance for varying numbers of clusters and cluster separation. After thresholding doublet scores based on the simulated doublet rate (5%), the recall (true positive rate) was measured using all doublets (*left*) or between-cluster doublets only (*right*). Error bars are standard deviation of 10 independent simulations. **(C)** Evaluation of doublet detector performance for varying cluster size asymmetry. Panels as in (B). Error bars are standard deviation of 10 independent simulations. Gray points correspond to individual simulations. **(D)** Evaluation of doublet detector performance for a branching continuum with varying branch separation. Recall was measured for all doublets (*left*) and when limiting to doublets formed by cells from opposite branches (*right*). Error bars and gray points as in (C). **(E)** Prediction of the detectable doublet fraction, *ϕD*, using the distribution of scores for the synthetic doublets. **(F)** Comparison of predicted *ϕD* to observed doublet recall for the simulations in (B).

The doublet detector also performed well when predicting doublets in a continuum of cell states: in a simulation of two paths diverging from the same starting state, up to 92% of doublets formed by cells from divergent states (>10% of the way towards opposite endpoints) were identified at a precision of 98% (**Fig. 2D**). As expected, doublets forming near the point of divergence were poorly identified. In summary, these results illustrate the basic concepts of the classifier in idealized settings with known inputs.

### Performance on dataset #1: human-mouse cell mixture

We tested the Scrublet on a publicly available dataset consisting of a mixture of human (HEK293 T) and mouse (NIH3T3) cells (**Fig. 3A**). This dataset, though not representative of most single-cell experiments, provides a useful test case because the differences between human and mouse genomic sequence provide an independent way to detect doublets [2, 3]. We defined a partial “ground truth” on doublet identity according to whether a cell barcode associates with transcripts from both species (a doublet), or just one species (**Fig. 3B**). Because doublets arising from the encapsulation of two human or two mouse cells cannot be identified as such, we expected our doublet detector to correctly predict all “ground truth” labeled doublets, since they arise from distinct human and mouse cell types.

**Figure 3.**
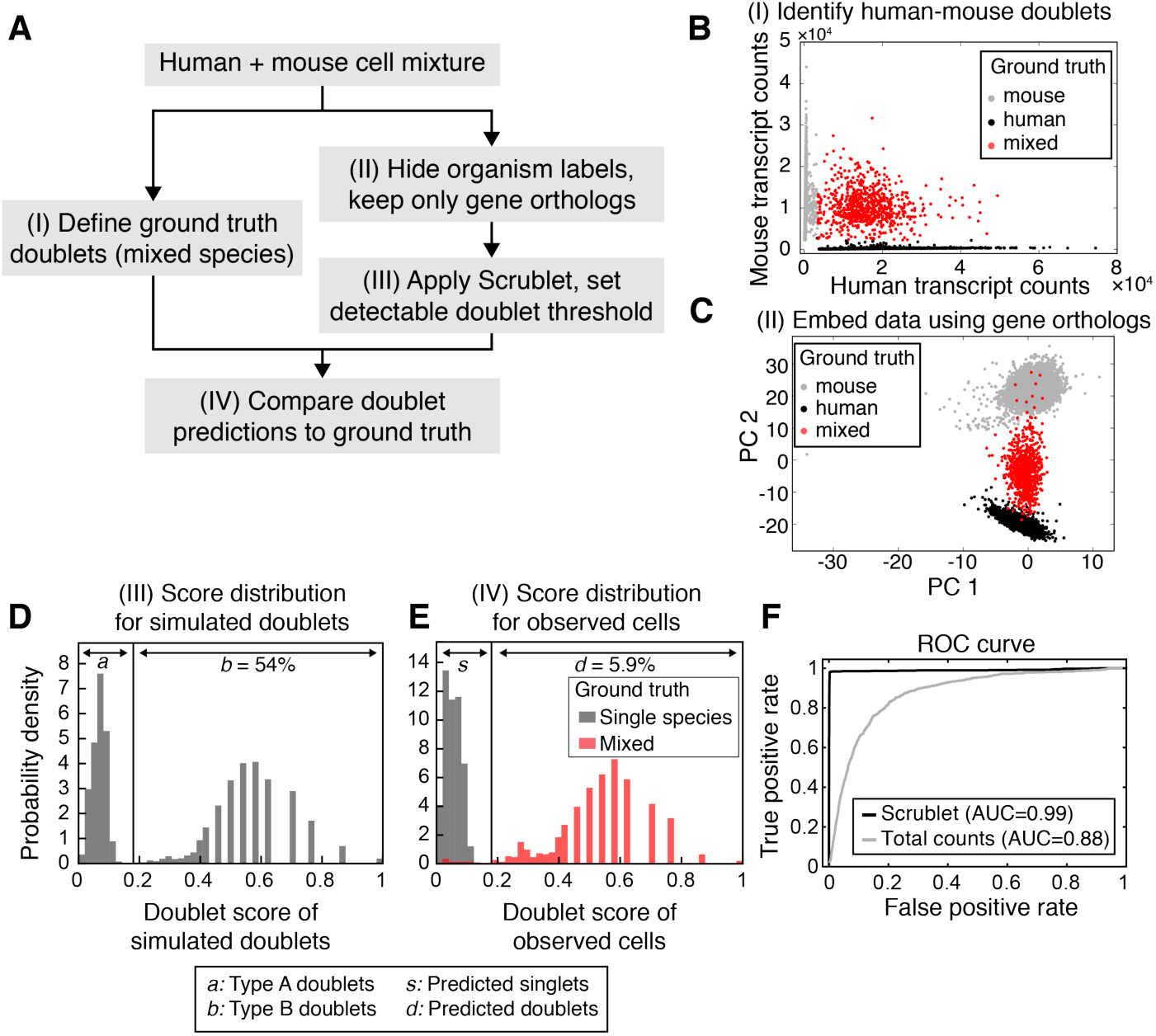
Doublet prediction for a mixture of human and mouse cells. **(A)** Schematic overview of species mixing experiment. **(B)** Identification of mixed-species doublets based on fraction of reads mapping to human or mouse transcriptome. **(C)** Principal component (PC) analysis of single-cell transcriptomes, restricting to human-mouse gene orthologs. **(D)** Histogram of doublet scores for simulated doublets. The bimodal distribution reflects the two types of doublets: undetectable intra-species Type A doublets (left peak) and inter-species Type B doublets (right peak). **(E)** Histograms of doublet scores for observed singlets (gray) and doublets (red). **(F)** Receiver-operator characteristic (ROC) curve for Scrublet and total transcript counts as predictors of inter-species doublets. AUC, area under the curve.

After hiding species labels and restricting to orthologous genes (**Fig. 3C**), Scrublet estimated the detectable (Type B) doublet fraction at *ϕD* = 54%, close to the 50% expected for cross-species doublets given equal input of mouse and human cells (**Fig. 3D**). Furthermore, the detector accurately identified human-mouse doublets with a receiver-operator characteristic (ROC) area under the curve (AUC) of 0.99 (recall of 98% of human-mouse doublets with precision of 96%) (**Fig. 3E,F**). In contrast, predicting doublets on the basis of total transcript counts was less effective (AUC=0.88), since the average human cell contained nearly twice as many transcripts as the average mouse cell; to achieve a recall of 90%, the precision dropped to just 15% (**Fig. 3F**).

### Performance on dataset #2: peripheral blood cells from multiple individuals

To test the doublet detector in a more typical experimental context, we evaluated its performance using a published dataset generated from a mixture of eight genotyped human donors’ mature blood cells (peripheral blood mononuclear cells, PBMCs) [11]. The authors identified “ground truth” multiplets as cell barcodes associated with reads containing polymorphisms from more than one individual (**Fig. 4A**). Given that the data represent a similar number of cells from each individual, roughly 7 out of 8 doublets should occur between individuals and can be identified using this approach. Thus, the “ground truth” is close to perfect, but 12.5% of true doublets are expected to be undetected.

**Figure 4.**
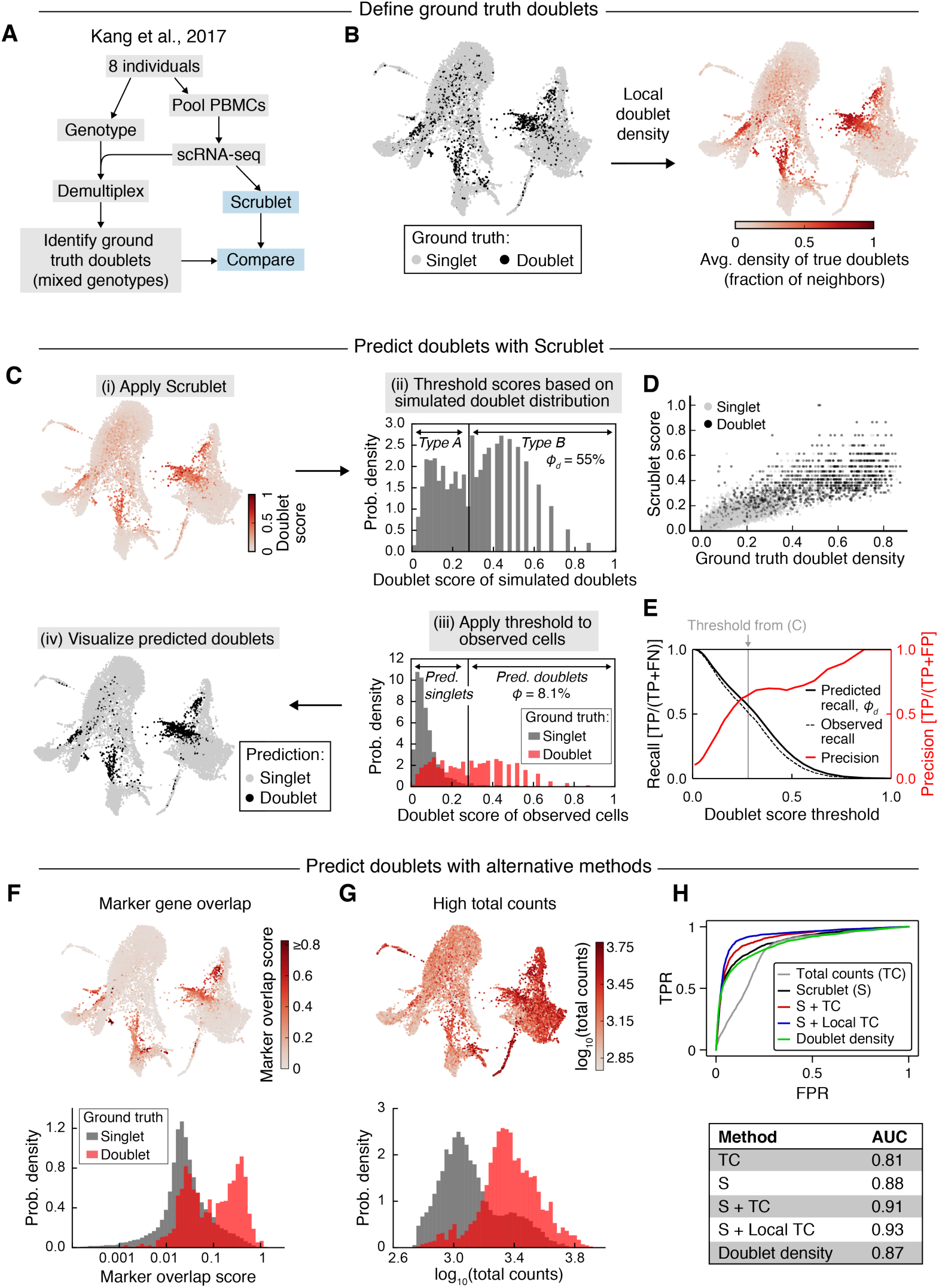
Doublet prediction for blood cells from eight genotyped human donors. **(A)** Schematic overview of genotyped cell mixing experiment. **(B)** *Left:* Force-directed graph layout of the profiled cells. Black points indicate ground truth doublets identified by demuxlet as barcodes associated with polymorphisms from more than one individual [11]. *Right:* Force-directed graph layout of ground truth doublet score, defined as the fraction of a cell’s neighbors that are mixed genotyped doublets. **(C)** Application of Scrublet to the transcriptomic data. After calculating doublet scores (i), the histogram of scores for simulated doublets was used to determine a threshold for detection of Type B doublets (ii). Applying this threshold to observed cell barcodes (iii) yielded doublet predictions for each transcriptome (iv). *ϕ_d_*, predicted detectable doublet rate; *ϕ*, fraction of transcriptomes predicted to be doublets. **(D)** Comparison of Scrublet to the ground truth doublet score, colored by genotype-based doublet labels (singlets, gray; doublets, black). **(E)** Comparison of detectable doublet fraction (solid black line) and actual recall (dashed black line) for a range of doublet score thresholds, and the corresponding precision (red line). TP, true positives; FN, false negatives; FP, false positives. **(F)** Alternative doublet prediction based on co-expression of marker genes of distinct cell types. *Upper:* force-directed graph layout with cells colored by marker overlap score. *Lower:* histograms of marker overlap score for ground truth singlets (gray) and doublets (red). **(G)** Alternative doublet prediction based total transcript counts. *Upper:* force-directed graph layout with cells colored by total counts. *Lower:* histograms of total counts for ground truth singlets (gray) and doublets (red). **(H)** ROC curves *(upper)* and AUC scores *(lower)* for various doublet prediction methods. “S+TC” and “S+Local TC” are linear combinations of the Scrublet score and total counts or the Scrublet score and total counts relative to neighboring cells, respectively (see **Extended Methods** for details).

To make use of this orthogonal method for multiplet detection, we compared Scrublet predictions to the ground truth doublets and also generated a ground truth score by calculating the fraction of each cell’s neighbors that were mixed genotype doublets (**Fig. 4B**). Because this score reflects the density of doublets in a region of gene expression space, it is directly comparable to the score computed using Scrublet. We then applied Scrublet to the transcriptomic data (**Fig. 4C**) and compared the Scrublet scores to these ground truth scores (**Fig. 4D**). This comparison showed a fair agreement: 89% of doublets with a high ground truth score (>0.4) were also identified by Scrublet, with a precision of 77%. The high-scoring cells for both methods co-localized in a low-dimensional visualization of the data (**Fig. 4B,C**), with undetected doublets scattered among other cell states. Furthermore, the recall of true doublets was accurately predicted by *ϕD*, the detectable doublet fraction as measured by the simulated doublet distribution (**Fig. 4E**). At the selected doublet score threshold, the recall of 49% was in good agreement with the *ϕD* of 55%, and this held true across a range of thresholds. This suggests that even though many doublets go undetected, the fraction of identifiable doublets can be accurately estimated. Though the precision was just 66%, this can be explained in part by the imperfect nature of the ground truth labels, since doublets formed by cells from the same individual are undetected.

As with the previous dataset, we compared the doublet detector performance to alternative strategies: (1) identifying cells co-expressing curated marker genes of distinct cell types, and (2) identifying cells with high total transcript counts. For the former method, we created a list of highly specific marker genes of each cell type in this dataset and then calculated the amount of co-expression of marker genes from different cell types (**Fig. 4 F**) to define a “marker co-expression score” (**Extended Methods**). Of the 773 true doublets correctly identified by Scrublet, 68% also had a high degree of marker gene co-expression. Overall, the “marker co-expression score” did not perform as well as Scrublet (AUC 0.77 vs. 0.88) and required significant manual annotation.

For the method relying on high total transcript counts, we found that true doublets did tend to have higher total transcript counts than singlets (AUC=0.81) (**Fig. 4G,H**). Because total counts appeared to be informative and did not require any manual annotation, we created a hybrid predictor by linear combination of each cell’s Scrublet score with its locally normalized total counts (**Extended Methods**). While this hybrid approach performed better than any other for this particular example (AUC=0.93) (**Fig. 4 H**), its effectiveness may vary across datasets, and it required additional parameter fitting.

### Performance on dataset #3: peripheral blood cells at multiple concentrations

In a third test, we turned to a dataset that offers a less direct independent strategy for detecting Type B doublets: namely, a single sample of PBMCs split and barcoded at two different cell concentrations, yielding either 4,352 (“PBMC-4 k”) or 8,391 (“PBMC-8 k”) transcriptomes. We reasoned that multiplet-specific cell states should be identifiable as clusters whose relative abundance increases with increasing input cell concentration, because in fully dissociated samples, a doubling of cell concentration doubles the probability of randomly encapsulating two cells into the same droplet. In the PBMC data, states comprised uniquely of doublets should double in relative abundance, with cell states that are predominantly singlets decreasing only incrementally (**Fig. 5A**).

**Figure 5.**
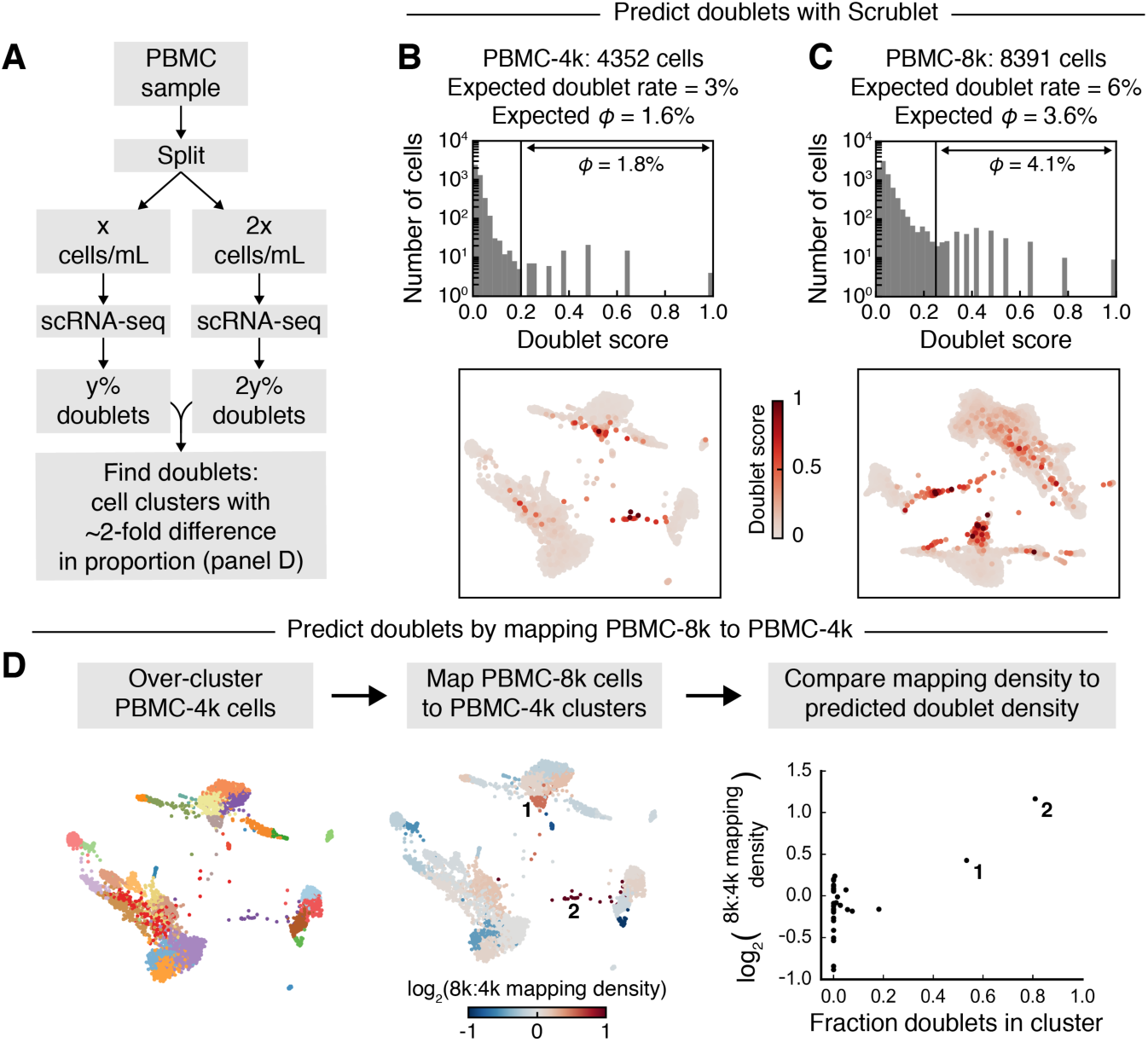
Doublet prediction using multiple concentrations of blood cells. **(A)** Schematic overview of how multiple concentrations of the same cell sample can be used to identify doublet-specific states. **(B)** Scrublet score histogram *(upper)* and force-directed graph layout *(lower)* for the low cell concentration (PBMC-4 k) sample. *ϕ*, fraction of transcriptomes predicted to be doublets. Expected *ϕ* was estimated using the expected doublet rate and the predicted detectable doublet fraction. **(C)** Same as (B), but for the high cell concentration (PBMC-8 k) sample. **(D)** Comparison of relative sizes of cell clusters in PBMC-4 k and PBMC-8 k samples to identify doublet-specific clusters. After clustering the PBMC-4 k cells *(left)*, each PBMC- 8 k cell was mapped to its most similar PBMC-4 k cell, and the proportions of cells from each sample in each cluster were compared *(center)*. This relative cluster abundance was then compared to the Scrublet predictions *(right)*.

As expected, the doublet detector identified roughly twice as many doublets in the PBMC-8 k sample (4.1%) as in PBMC-4 k (1.8%) (**Fig 5B,C**). Furthermore, when we compared the PBMC-8 k cells to their most similar PBMC-4 k counterparts, the predicted doublet states were present at a higher relative abundance, while singlet states changed little or decreased (**Fig. 5D**). This test again suggests that the doublet detector correctly identifies Type B doublets.

### Prediction of doublets in a cell state continuum

The above examples demonstrate Scrublet’s ability to correctly identify Type B doublets from datasets consisting of distinct cell types. In a final example, we applied it to a continuum of cell states by analyzing transcriptomes of Kit+ hematopoietic progenitors from the mouse bone marrow [19] (**Fig. 6A**). These cells form a continuum from multipotent progenitors to unilineage committed cells. Several groups of doublets were readily distinguishable (**Fig. 6B–C**) and formed “bridges” between different committed progenitor types. Here we lack a ground truth for confirming the identity of the doublets, but since such bridges are inconsistent with our current understanding of hematopoiesis, it is likely that our doublet detector is correct in identifying them.

**Figure 6.**
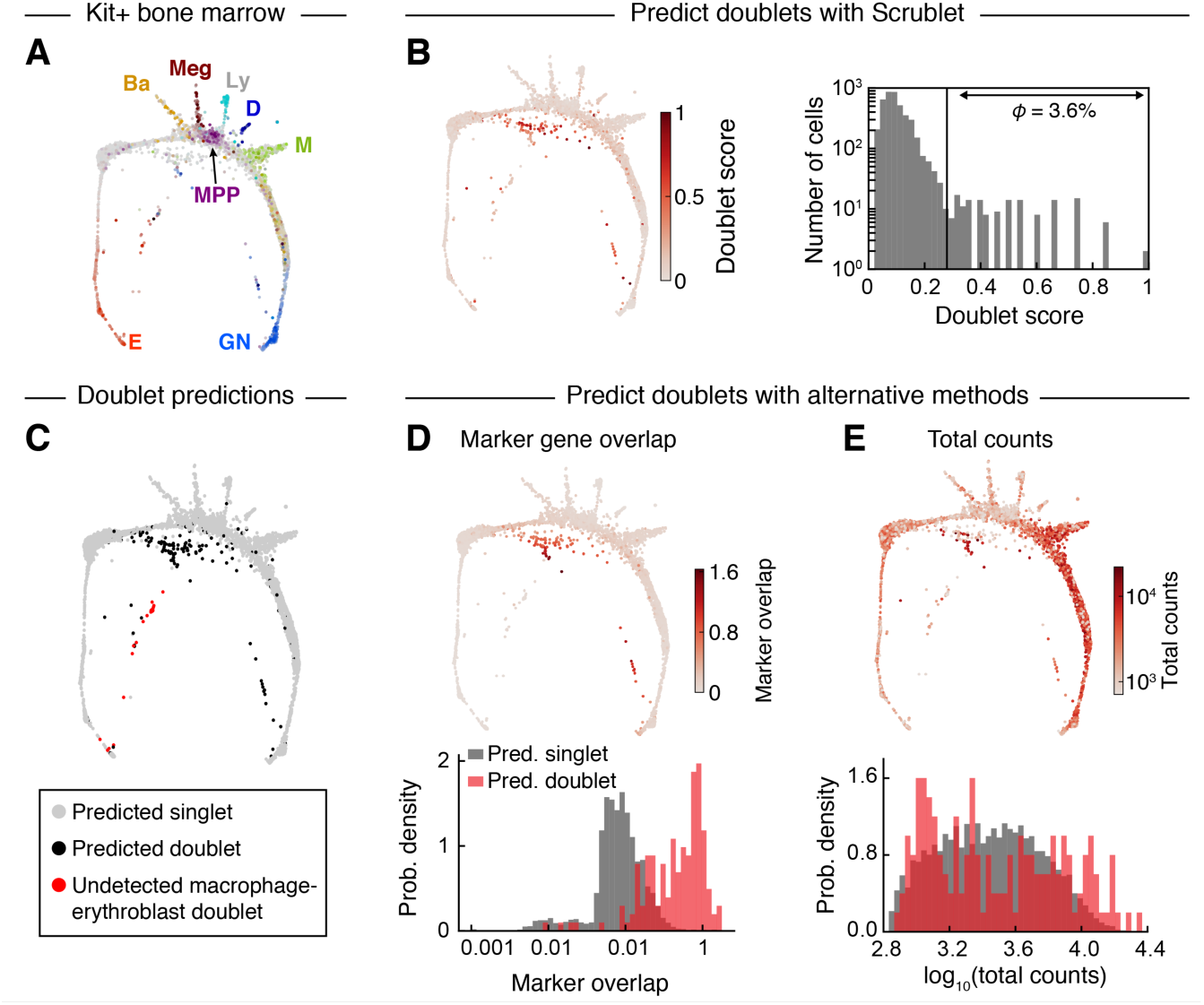
Prediction of doublets in a continuum of differentiating hematopoietic progenitors. **(A)** Force-directed graph layout of Kit+ mouse bone marrow cells profiled by scRNA-seq. Cells are colored by expression of established marker genes. E, erythroid; Ba, basophil/mast cell; Meg, megakaryocyte; MPP, multipotent progenitor; Ly, lymphoid; D, dendritic cell; M, monocyte; GN, granulocytic neutrophil. Adapted from [19]. **(B)** Force-directed graph layout colored by Scrublet score *(left)* and histogram of Scrublet scores *(right). ϕ*, fraction of transcriptomes predicted to be doublets. **(C)** Predicted doublets localized on force-directed graph layout. Gray, predicted singlets; black, Scrublet-predicted doublets; red, likely erythroblast-macrophage doublets (*C1qa*+ *Hba-a1*+), undetected by Scrublet due to absence of macrophage singlets in the Kit+ data. **(D)** Alternative doublet prediction based on co-expression of marker genes of distinct cell types. *Upper:* force-directed graph layout with cells colored by marker overlap score. *Lower:* histograms of marker overlap score for Scrublet-predicted singlets (gray) and doublets (red). **(E)** Alternative doublet prediction based on total transcript counts. *Upper:* force-directed graph layout with cells colored by total counts. *Lower:* histograms of total counts score for Scrublet-predicted singlets (gray) and doublets (red).

We again compared Scrublet to other approaches based on marker genes or total counts. As before, predicted doublets consistently expressed combinations of marker genes for distinct maturing progenitor states (**Fig. 6D**), while only some predicted doublets had above average total transcript counts (**Fig. 6E**).

This dataset was also instructive in highlighting a shortcoming of our approach when one of its assumptions is violated. Namely, Scrublet can detect cell aggregate doublets only if both parent cell types are observed as singlets elsewhere in the dataset. Through manual curation, we identified a small group of transcriptomes co-expressing markers of erythroblasts and mature macrophages (**Fig. 6D**). Macrophages and Kit+ erythroblasts are known to physically associate in the bone marrow in erythroblastic islands [20] and have been observed in other scRNA-seq datasets [21]. Since macrophages do not express the cell surface receptor Kit, used for cell purification in this experiment, they appear only in the form of doublets in this dataset. Unfortunately, such aggregates might confound other methods for doublet detection, including all of the experimental methods in **Table 1**. They may, however, be identifiable by combining multiple datasets in order to provide the full set of singlet states for Scrublet.

## Discussion

We proposed and tested a classification scheme for cell multiplets, focusing on cell doublets, as these are expected to form the majority of multiplets in all but specialized cases. The classifier is trained using the data itself and reasonable assumptions about the structure of gene expression space. The application to simulated data, and then to four empirical datasets, demonstrates that the approach can accurately identify doublets formed by cells from distinct states, as assessed by formal estimates of recall and precision where possible. The cell transcriptomes that scored as doublets with highest confidence were also those found after manual curation of the data to co-express marker genes of distinct cell states. The classification approach outperformed manual curation and simple total counts-based approaches, although it benefitted from being combined with total counts information.

Although the method appears to perform well, its underlying assumptions do impose some limitations. First, the method assumes that multiplets are rare. This is required (1) to justify the study of doublets rather than all multiplets and (2) for the doublets simulated by the classifier to overwhelmingly reflect doublet states rather than higher-order states. Second, the method strictly requires that every cell state contributing to a doublet also be represented as a singlet state in the dataset. If a particular singlet cell state is excluded experimentally, it trivially cannot be detected as part of a multiplet state, because the missing parent state does not contribute to the simulated doublet pool used for doublet classification. This limitation could be appreciated in the final dataset, from cells purified conditional on expression of a cell surface protein, Kit. We found that cell doublets resulting from incomplete dissociation of a Kit+ erythroid cell and a Kit- macrophage could not be detected by the classifier, because no singlet macrophage state was present in the dataset. An extension of this limitation is that the method could underperform if cell clumps with a stereotyped composition occur in a sample. Scrublet performs best for doublets resulting from random co-encapsulation because the frequency of such doublet states can be predicted by the frequency of singlet states. Doublets from incomplete dissociation can still be effectively detected provided that they are rare and that the singlet states are well represented in the data.

A third limitation of the approach is its sensitivity to the structure of the single-cell state manifold. Scrublet performs best in identifying doublets formed between distinct parent states. This limitation is quantified for any given dataset by the calculated detectable doublet fraction, *ϕD*, which is expected to be high if singlet states are distributed among many discrete, well-separated states; it is only 50% if cells form two discrete and equal-sized clusters, and it can be lower than 50% for complex continuum manifolds. Countering this shortcoming is the notion that rare doublet states are only important to exclude if they form novel features on a cell state manifold, which would in turn make them detectable using the proposed approach. Therefore, the doublet detector provides a useful tool for both estimating the potential impact of doublets on downstream hypothesis generation through the magnitude of *ϕD*, and for identifying bona fide doublet states for exclusion.

## Availability

Python code and examples implementing the doublet detector are provided at github.com/AllonKleinLab/scrublet. Scrublet has also been incorporated into SPRING (kleintools.hms.harvard.edu/tools/spring.html), an interactive tool for single-cell data exploration [22].

## Acknowledgments

This work was supported by NIH grants 1R01HL141402 and 5R33CA212697 and an Edward Mallinckrodt, Jr. Foundation Grant. We thank James Briggs for insightful discussion in conceptualizing the approach.

## Author Contributions

SLW conceived the approach. SLW, RL, and AMK formalized the problem. SLW and RL developed and applied the methods. SLW and AMK wrote the paper. AMK supervised the work.

## Extended Methods

### The Scrublet algorithm

#### General approach

Starting with a raw counts matrix, *X*, where *X_i,j_* is the number of detected transcripts of gene *j* in cell *i*,

1. Pre-filter cell barcodes to exclude background, typically barcodes with insufficient total transcripts detected.
2. Simulate doublets by combining the counts from random pairs of cells: the counts for gene *j* in doublet *i*^′^ with parent cells *a* and *b* is *X_i^′^j_*, = *X_a,j_* + *X_b,j_*.
3. Build a k-nearest-neighbor (kNN) classifier, labeling observed cells as 0 and simulated doublets as 1. In detail, construct a kNN graph using the union of observed cells and simulated doublets and calculate the doublet score as the fraction of neighbors that are simulated doublets.
4. Remove likely doublets by thresholding the doublet scores or by clustering observed cells and identifying clusters with uniformly high scores.

#### Detailed method

Throughout this paper, and in the code provided online at github.com/AllonKleinLab/scrublet, we implement the above approach as follows.

##### Preprocessing

Starting with a background-filtered, UMI-based counts matrix for the observed cells, we perform normalization, gene filtering, and principal components analysis (PCA):

1. Normalize each cell by its total counts, setting the post-normalization total to the average total of all cells.
2. Identify highly variable genes, keeping genes with ≥ *n_g_* counts in ≥ *n_c_* cells and in the top *q^th^* percentile of most variable genes, as measured by *V*-score (baseline-corrected Fano factor) [2].
3. Z-score normalize at the gene level.
4. Run PCA.

##### Doublet simulation

Because PCA is a linear transformation, we simulate doublets by averaging the PCA coordinates of the randomly sampled parent cells, weighting by the total transcripts in each parent. That is, if doublet *i^′^* is generated by parent cells *a* and *b* with transcript count totals *t_a_* and *t_b_* and PCA coordinates *P_a_* and *P_b_*, then the PCA coordinate for doublet *i*^′^ is 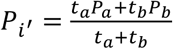

##### kNN classifier

Following PCA, a kNN graph is built using Euclidean distance in the combined PCA embedding of the observed and simulated cells. Because both the number of neighbors, *k*, and the ratio of the number of simulated doublets to observed cells, *r*, are user-provided parameters, *k* is scaled by *r*, and the adjusted number of neighbors, *k_adj_* = round(*k* ⋅ 1 + *r*), is used to construct the graph.

Next, *f_i_* and *f_i_^′^*, the doublet scores for the observed cells *i* and simulated doublets *i^′^*, respectively, are calculated by finding the fraction of each cell’s (or simulated doublet’s) neighbors that are simulated doublets, adjusting for *r* accordingly. We also rescale the scores by the expected doublet rate *d*, though this information is not essential for obtaining an interpretable result (a default of *d* = 0.1 is reasonable; see below for additional details). For cell *i* with *n_i_* observed cell neighbors and *m_i_* simulated doublet neighbors, the doublet score is

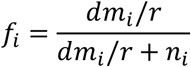

and similarly for *f_i_^′^*.

##### Setting the doublet score threshold

After computing the doublet scores *f_i_* and *f_i_^′^*, a threshold *θ* is set based on the distribution of *f_i_^′^*, and observed transcriptomes with *f_i_* < *θ* as predicted as doublets. In all of the presented examples, the distribution of *f_i_^′^* was bimodal, reflecting the differences between Type A and Type B doublets, and the threshold was set by eye to lie between the two peaks of the histogram of *f_i_^′^*.

##### The role of the expected doublet rate

While the expected doublet rate *d* does not directly influence the doublet predictions, it does play a role in two ways:

1. Rescaling the doublet scores: setting *d* near the true doublet rate results in a more bimodal distribution of *f_i_^′^* and, similarly, better separation of the observed doublet scores.
2. “Sanity checking” predictions: after setting the threshold *θ*, the value of *d* can be compared to the resulting predicted doublet rate. If *ϕ* is the fraction of observed transcriptomes with *f_i_* < *θ* and *ϕ_D_* is the fraction of simulated doublets with *f_i_^′^* < *θ*, then the predicted overall doublet rate is *ϕ/ϕ_D_*. This predicted doublet rate should roughly agree with *d*.

### Testing Scrublet

#### Splatter simulations

We used the Splatter R package (v1.0.3) [18] to simulate ground truth data for testing the doublet detector. For each set of parameters, we simulated 10 replicates with 5000 cells and 2000 genes, using default parameters except where noted below. Doublets were simulated at a rate of 5% by randomly sampling (without replacement) pairs of cells and summing their counts; cells used to generate doublets were then removed from the data. **Table 2** summarizes the conditions simulated for **Fig. 2**.

**Table 2.**

| Panel | Number of groups | Group1 size / Group2 size | Splatter parameter “method” | Splatter parameter “mean.shape” | Splatter parameter “de.prob” |
| --- | --- | --- | --- | --- | --- |
| B | 2, 3, 5, 10, 15 | n/a (all uniform) | groups | 0.5 | 0.005, 0.01, 0.02, 0.03, 0.04, 0.05, 0.1 |
| C | 2 | 1, 1.5, 2.3, 4, 9, 19 | groups | 0.5 | 0.05 |
| D | 2 | 1 | path | 0.5 | 0.01, 0.02, 0.05, 0.1, 0.15, 0.25, 0.4 |

Prior to predicting doublets, PCA was run using genes with at least 3 counts in at least 3 cells. For **Fig. 2B,C**, we used all PCs with eigenvalues that were at least 20% of the maximum eigenvalue. For **Fig. 2D**, the top 4 PCs were used for all conditions. The doublet detector was run using *k* = 40, *r* = 5, and *d* = 0.05.

To determine the overall recall 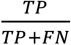; TP, true positives; FN, false negatives), we set a doublet score threshold based on the simulated doublet rate of 5%; that is, cells with doublet scores in 95th percentile or above were labeled as predicted doublets. Thus, the precision (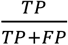; FP, false positives) is equal to the recall. The same procedure was used to measure the recall for between-cluster doublets, restricting to doublets formed by cells from different groups. For the branching continuum simulation, between-branch doublets were defined as doublets formed by cells on opposite branches and with Splatter pseudotime >10%.

#### Human-mouse dataset

##### Pre-processing and doublet detector parameters

Separate pre-filtered counts matrices for human and mouse genes were downloaded from 10X Genomics (support.10xgenomics.com/single-cell-gene-expression/datasets/2.1.0/hgmm_12 k), along with species assignments for each barcode (6,164 human cells, 5,915 mouse cells, and 741 mixed human/mouse multiplets). To create a single counts matrix blind to the species of origin, each cell’s UMI counts for genes with identical mouse and human names (*n* = 15,642 genes) were added together, and all other genes were excluded. For PCA, we used the top 20% most highly variable genes with ≥3 counts in ≥5 cells (*n*=2,372 genes) and kept the first two PCs. Scrublet was run using *k* = 50, *r* = 10, and *d* = 0.12 (twice the observed rate of human-mouse doublets). To classify cells as singlets or doublets, a threshold was set by eye using the histogram of doublet scores for simulated doublets (**Fig. 3D**).

#### Demuxlet PBMC dataset

##### Pre-processing and doublet detector parameters

A filtered counts matrix (14,619 cells and 35,635 genes) was downloaded from GEO (accession ID GSM2560248), and demuxlet singlet/doublet calls were obtained from the paper’s GitHub page (github.com/yelabucsf/demuxlet_paper_code). For PCA, we used the top 25% most highly variable genes with ≥2 counts in ≥3 cells (*n*=3,197 genes) and kept the first 25 PCs. Scrublet was run using *k* = 50, *r* = 5, and *d* = 0.11 (the observed doublet rate). To classify cells as singlets or doublets, a threshold was set by eye using the histogram of doublet scores for simulated doublets (**Fig. 4C**).

##### Ground truth doublet score

The ground truth doublet score was created by building a kNN graph (*k* = 35) using the observed cells and calculating the fraction of each cell’s neighbors labeled as doublets by demuxlet.

##### 2-D visualization

Transcriptomes were visualized using a force-directed layout of the four-nearest-neighbor graph of observed cells, where neighbors were identified using Euclidean distance in PC space.

##### Marker gene co-expression score

The marker gene co-expression score was created by identifying highly specific marker genes for each cell type, smoothing expression of these genes over the four-nearest-neighbor graph (see “Graph-based smoothing”, below), and summing the products of pairs of non-overlapping marker genes. In detail, we combined the following pairs of marker genes:

- T-cell and NK cell: *CD27* x *SH2D1B, CD27* x *IGFBP7, CD27* x *KLRF1*
- T-cell and B-cell: *CD27* x *BANK1, CD27* x *BLK, CD27* x *MS4A1*
- T-cell and monocyte: *CD27* x *CST3*
- B-cell and NK cell: *BANK1* x *SH2D1B*

Letting 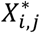be the smoothed, normalized gene expression of gene *j* in cell *i*, the composite score for a pair of genes *a* and *b* is

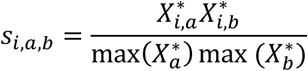

For a given cell type pair *p* with gene pairs 1,2, …, *n*, the marker gene overlap score for cell *i* is defined as

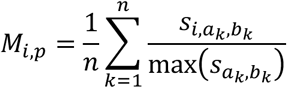

And the composite marker gene overlap score for all cell type combinations (as shown in **Fig. 4F**) is *Σ_p_ M_i,p_*.

##### Hybrid doublet score (Scrublet + total counts)

For this dataset, we also tested whether combining total counts information with the Scrublet score would improve doublet classification, e.g., by enabling detection of Type A doublets (**Fig. 4H**). In both versions described below, the parameters (relative weights of Scrublet and total counts-based scores) were fit to maximize the AUC.

1. We tested a simple linear combination of Scrublet (*f_i_*) and total counts (*T_i_*): 4*f_i_* + log_10_(*T_i_*).
2. We created a “local total counts” (*L_i_*) score, defined as a cell’s total counts divided by the average total counts of its simulated doublet neighbors, and combined it with Scrublet: 3*f_i_* + *L_i_*.

#### PBMCs at multiple concentrations

##### Pre-processing and doublet detector parameters

Filtered counts matrices were downloaded from 10X Genomics (PBMC-4 k: support.10xgenomics.com/single-cell-gene-expression/datasets/2.1.0/pbmc4 k; PBMC-8 k: support.10xgenomics.com/single-cell-gene-expression/datasets/2.1.0/pbmc8 k). For PCA, we used the top 15% most highly variable genes with ≥3 counts in ≥3 cells (PBMC-4 k, *n*=1,129 genes; PBMC-8 k, *n*=1,307 genes) and kept the first 30 PCs. Scrublet was run using *k* = 50, *r* = 5, and *d* = 0.03 (PBMC-4 k) or *d* = 0.06 (PBMC-8 k), based on the expected doublet rates (support.10xgenomics.com/permalink/3vzDu3zQjY0o2AqkkkI4CC). To classify cells as singlets or doublets, a threshold was set by eye using the histogram of doublet scores for simulated doublets.

##### 2-D visualization

Transcriptomes were visualized using a force-directed layout of the four-nearest-neighbor graph of observed cells, where neighbors were identified using Euclidean distance in PC space.

##### Mapping PBMC-8 k to PBMC-4 k

To map the PBMC-8 k data to the PBMC-4 k data, we TPM-normalized both datasets, ran PCA on the PBMC-4 k cells, and used the same eigenvectors to transform the PBMC-8 k data. The PBMC-4 k cells were clustered using spectral clustering of the four-nearest-neighbor graph with 30 clusters. We then mapped each PBMC-8 k cell to its nearest PBMC-4 k cell (Euclidean distance) and calculated the number of PBMC-8 k cells mapping to each PBMC-4 k cluster. In **Fig. 5D**, we present the relative number of PBMC-8 k cells per cluster; that is, if *n_j_* is the number of PBMC-8 k cells mapping to cluster *j* and *N_4k_* and *N_8k_* are the total number of PBMC-4 k and PBMC-8 k cells, then the relative mapping frequency for cluster *j* is 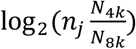.

#### Hematopoietic progenitor dataset

##### Pre-processing and doublet detector parameters

The raw counts matrix was downloaded from GEO (GSM2388072). Restricting to cells from library batches 2, 3, and 4, we also excluded cells with fewer than 700 total counts or with >15% mitochondrial gene counts (n=4,273 cells final). For PCA, we filtered genes using the same method as the original paper [19], keeping genes with mean expression >0.05 counts and a coefficient of variation >2 (*n*=7,255 genes), and kept the first 40 PCs. Scrublet was run using *k* = 50, *r* = 5, and *d* = 0.1. To classify cells as singlets or doublets, a threshold was set by eye using the histogram of doublet scores for simulated doublets. After removing high-scoring cells (Scrublet score >0.28, n=146 cells), we re-ran Scrublet and observed additional likely doublets that had been residing at the core of a dense doublet cluster in the original data (round 2 Scrublet score >0.28, n=34 cells). Following removal of these cells, a third round of Scrublet yielded no additional likely doublets.

##### 2-D visualization

Transcriptomes were visualized using the force-directed graph layout appearing in the original publication, with minor modifications. Because the published plot was generated after removing doublets, we added doublets back to the visualization by building a kNN graph (k=4) with all transcriptomes (filtered as described above) and running a force-directed graph layout with the positions of the original cells fixed in place, allowing the remaining cells to relax.

##### Marker gene co-expression score

The marker gene co-expression score was created by identifying highly specific marker genes for each cell type, smoothing expression of these genes over the four-nearest-neighbor graph (see “Graph-based smoothing”, below), and summing the products of pairs of non-overlapping marker genes. The combined marker overlap score was calculated as described in the “Demuxlet PBMC dataset” section, above.

We combined the following pairs of marker genes to identify doublets that were also detected by Scrublet (**Fig. 6D**):

- Early erythroid and early neutrophil: *Car1* x *Mpo*
- Early erythroid and late neutrophil: *Car1* x *Ngp*
- MPP and late neutrophil: *Cd34* x *Ngp*

And to identify macrophage-erythroblast doublets (n=37 cells) undetected by Scrublet (**Fig. 6C**):

- *C1qa* x *Hba-a1*

#### Graph-based smoothing

We used a diffusion-based method to smooth data over the kNN graph for the purposes of finding overlapping marker gene expression (**Figs. 4 F,6D**). In detail, we computed the smoothing operator *S = expm(-βL)*, where *L* is the Laplacian matrix of the kNN graph, *β* is the strength of smoothing (*β* = 1 throughout), and *expm* is matrix exponential (`scipy.linalg.expm from the SciPy Python package`). If *X^*^* is the smoothed version of gene expression vector *X*, then *X^*^* = *SX*.

